# Sediment source and dose influence the larval performance of the threatened coral *Orbicella faveolata*

**DOI:** 10.1101/2023.09.22.559035

**Authors:** Xaymara M. Serrano, Stephanie M. Rosales, Margaret W. Miller, Ana M. Palacio-Castro, Olivia M. Williamson, Andrea Gomez, Andrew C. Baker

**Affiliations:** Cooperative Institute of Marine and Atmospheric Studies, Rosenstiel School of Marine, Atmospheric, and Earth Science, University of Miami, Miami, Florida, USA; National Oceanic and Atmospheric Administration, Atlantic and Oceanographic Meteorological Laboratory, Miami, Florida, USA; SECORE International, Hilliard, Ohio, USA; Department of Marine Biology and Ecology, Rosenstiel School of Marine, Atmospheric, and Earth Science, University of Miami, Miami, Florida, USA; National Oceanic and Atmospheric Administration, Greater Atlantic Regional Fisheries Office, Gloucester, Massachusetts, USA

## Abstract

The effects of turbidity and sedimentation stress on early life stages of corals are poorly understood, particularly in Atlantic species. Dredging operations, beach nourishment, and other coastal construction activities can increase sedimentation and turbidity in nearby coral reef habitats and have the potential to negatively affect coral larval development and metamorphosis, reducing sexual reproduction success. In this study, we investigated the performance of larvae of the threatened Caribbean coral species *Orbicella faveolata* exposed to sediments collected from a reef site in southeast Florida recently impacted by dredging (Port Miami), and compared it to the performance of larvae exposed to sediments collected from the offshore, natal reef of the parent colonies. In a laboratory experiment, we tested whether low and high doses of each of these sediment types affected the survival, settlement, and respiration of coral larvae compared to a no-sediment control treatment. In addition, we analyzed the sediments used in the experiments with 16S rRNA gene amplicon sequencing to assess differences in the microbial communities present in the Port versus Reef sediments, and their potential impact on coral performance. Overall, *O. faveolata* larvae exposed to high doses of either sediment type (Port or Reef) exhibited reduced survival and settlement rates, but only the Port sediments resulted in adverse effects in the low-dose treatment. Sediments collected near the Port also contained different microbiomes than Reef sediments, and higher relative abundances of the bacteria Desulfobacterales, which has been associated with coral disease. We hypothesize that differences in microbiomes between the two sediments may be a contributing factor in explaining the observed differences in larval performance. Together, these results suggest that the settlement success and survival of *O. faveolata* larvae are more readily compromised by encountering port inlet sediments compared to reef sediments, with potentially important consequences for the recruitment success of this species in affected areas.

## Introduction

Dredging and port construction activities can induce chronic sedimentation and turbidity stress in surrounding coral reef habitats [1–6]. Sedimentation caused by dredging is often more harmful to corals than sedimentation caused by other activities, due to its acute onset and often long duration, typically lasting many months to years [1, 7]. Sedimentation effects vary depending on grain size and composition (e.g., [7–12]), and dredging activities can release large quantities of sediments in the water column that are typically of finer grain size compared to naturally occurring sediments (<63 μm, [7]). These finer sediments tend to attenuate more light and are more prone to being resuspended, thus causing a greater net reduction in light essential for photosynthesis [10]. Finer sediments also have an adhesive, clay-like texture that is more resistant to bioturbation when deposited [13], can lead to a change in the bacterial community [12], and is more likely to become anoxic [8, 14]. Finally, suspended sediment plumes, especially those with finer grain size, can also travel several km away from the dredging site (e.g., [9, 15, 16]), and can also release contaminants [17–19] and pathogens [11, 20, 21] which may potentially transmit coral diseases such as Stony Coral Tissue Loss Disease [22].

Sediments can negatively affect corals throughout their life cycle, suppressing coral health, condition, and survival via multiple mechanisms (reviewed in [7, 12, 13, 23]). Reported effects on early life stages of corals include reductions in the fecundity of parent colonies [24], fertilization [25, 26], larval settlement [27–31], and recruit survival [5, 27, 32, 33]. Tolerance to sedimentation exposure is estimated to be an order of magnitude lower for coral recruits compared to adults [13, 23, 34], although most of this work has been conducted on Pacific coral species (reviewed in [12, 13, 23]). Much less is known regarding the effects of sediments on coral larvae and recruits for Atlantic coral species, and this information is critical for developing appropriate conservation plans for these species. Poor understanding of coral responses to sediment disturbances during their most susceptible stages can result in inappropriate management of dredging projects, lead to preventable coral losses, and unnecessarily high costs to adaptively manage operations (corrective actions) and provide compensatory mitigation to offset impacts.

Sediments may interfere with the ability of coral larvae to find a suitable substrate on which to settle, which is key to successful recruitment and survival [31, 35, 36]. Even a thin layer of sediments not harmful to adult corals can be detrimental to coral larvae [23, 28, 31, 37–39], both by physically obstructing settlement surfaces [27, 30], and impairing biotic settlement cues such as those from crustose coralline algae (CCA) [30]. However, the mechanisms and cues by which sediments can affect settlement and recruitment success, or benthic habitat quality (e.g., [31]) are poorly understood. Recent work suggests that coral larvae show a preference for distinct microbial and/or chemical signature types in certain types of CCA [40–43], suggesting that specific bacterial communities can serve as environmental cues for coral settlement [40, 42]. How and whether microbial communities present in sediments can negatively affect the ability of coral larvae to identify these cues remains an open question.

The aim of the present study was to investigate the performance of coral larvae of an important Caribbean reef-building species (*Orbicella faveolata*) following a short-term (pulse) sediment exposure. *O. faveolata* (Ellis and Solander, 1786) is listed as threatened under the U.S. Endangered Species Act [44] but is present in southeast Florida in the vicinity of two large ports with expansions (deepening/widening) planned in the near future (Port Everglades and Port Miami). The most recent large-scale deepening project at Port Miami (which occurred from 2013-2015) resulted in severe sedimentation impacts to local coral reef habitats [1, 3, 15, 45], although the severity of those impacts has been disputed [46]. In addition, recent work exposed adult *O. faveolata* corals to Port Miami sediments during a 96-hr period and showed adverse effects on tissue regeneration capacity compared to no sediment controls [47]. In this study, we exposed *O. faveolata* larvae to sediments collected near the Port Miami channel and compared their performance with larvae exposed to reef sediments collected from the natal reef of the parent colonies in the Florida Keys. We compared the effects of the two different sediment types on the survival, settlement, and respiration of coral larvae, and also characterized the bacterial communities of these sediments using 16S rRNA gene amplicon sequencing to assess the potential role of microbial communities in affecting coral performance. To our knowledge, this is the first study to investigate the effects of suspended sediments on larvae of western Atlantic (Caribbean) coral species, and the first to use sediments that have not been dried or sterilized prior to exposure, which may have masked some of the biological or chemical effects of sedimentation in previous studies.

## Methods

### Coral larvae collection

*O. faveolata* larvae were collected and cultured as described in [48] and [49]. Gamete bundles were collected seven nights after the full moon (night of 14^th^ August 2017) from colonies that spawned at two reefs in the upper Florida Keys (Horseshoe and Sand Island reefs). Once the bundles broke apart, equal volumes of gametes from 3-4 parental genets were combined for fertilization. Resulting embryos were maintained in water collected from their natal reef at a land-based facility in the Keys (either in static bins with regular water changes or in a recirculating system with mesh-floored bins and a constant drip input) and cultured for ∼2 days so that unfertilized eggs would disintegrate, leaving behind only developing larvae. Embryos were kept in a shaded area under ambient light conditions and a mean temperature of 29°C. Larvae were then transferred to the University of Miami for experiments.

### Sediment collections and maintenance

Immediately prior to starting the larval experiments (24 - 48 h), two batches of sediments were collected for experiments, one from a “Reef” origin and another from a “Port” origin. Reef sediments were collected at Horseshoe Reef in Key Largo, from the same site where the coral parent colonies occurred (25°8’19”N, 80°17’41”W, Fig 1). Port sediments were collected from a site located ∼200 m north of the recently-dredged channel of Port Miami (25°45’50”N, 80°5’56”W, Fig 1), reported in [3] as having detrimental effects from dredging-induced sedimentation on adult corals. This site was shown by [3] as the most severely impacted location surveyed, with the highest sediment cover (43%) and highest prevalence of sediment “halos” (indicating partial mortality) compared to the reference stations.

**Fig 1.**
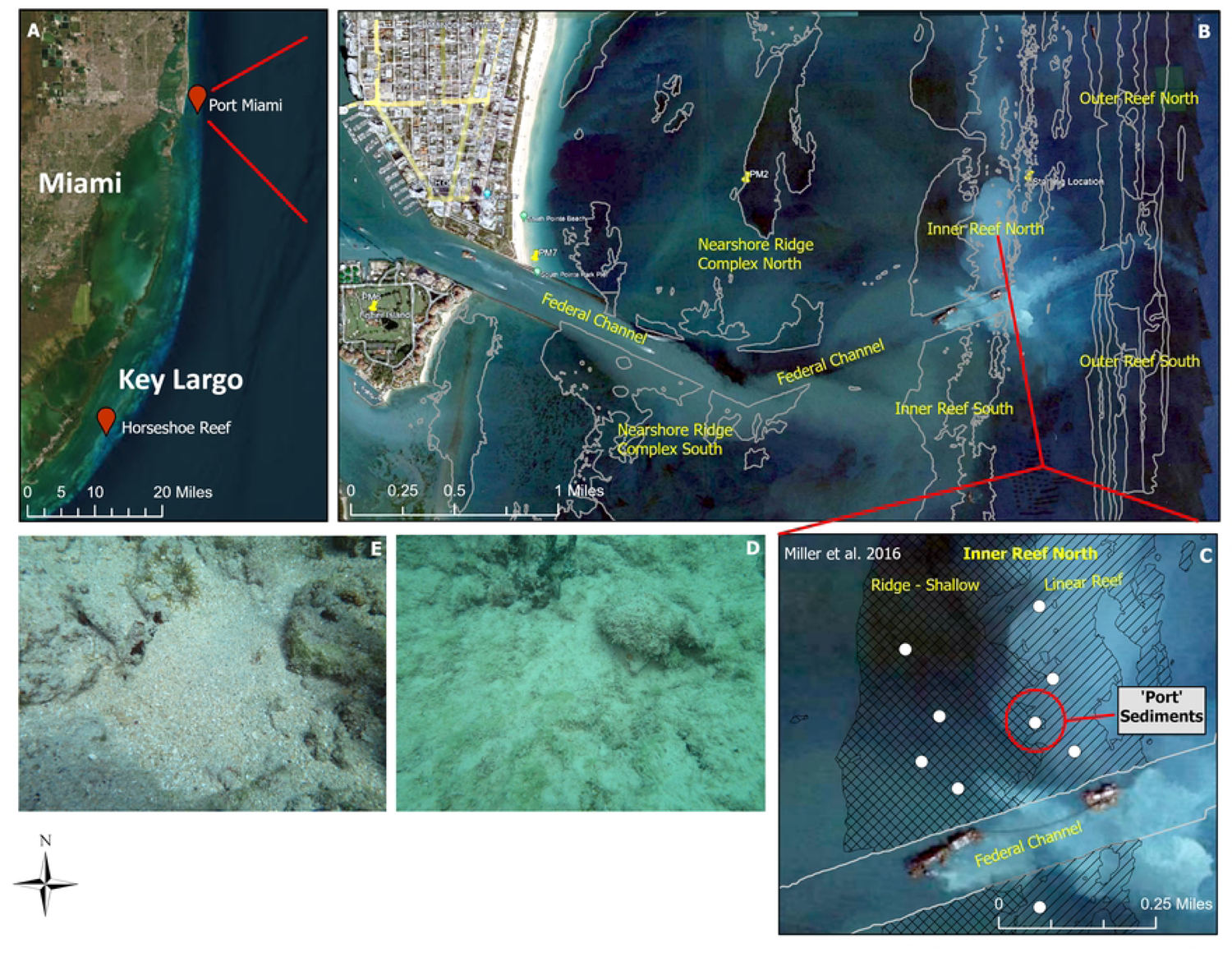
(A) Map of locations on the coast of southeastern Florida where Port and Reef sediments were collected from (near Port Miami and Horseshoe reef in Key Largo). (B) Map of coral reef habitats around the Port of Miami federal channel. (C) Port sediments were collected from a site within the Inner Reef located ∼200 m north of the recently dredged channel of Port Miami (circled in red), which was the most severely impacted location surveyed due to sedimentation compared to the reference stations by [3]. (D) Representative photo of the sediments collected near the Port of Miami. (E) Representative photo of the sediments collected at Horseshoe reef.

All sediments were taken from the upper ∼3 cm of the sediment layer (surficial layer) by sliding a metal tool below the sediment-water interface and then scooping them into a certified-clean HPDE sample container. Three to four containers were collected per site and combined to produce a single “batch” for experiments. Sediments were then brought to the laboratory, homogenized in a beaker of seawater and kept in continuous movement and aeration until used in experiments. Prior to use, sediments were allowed to settle and water was siphoned out with care to avoid disturbing the fine particles on the surface. Sediments were then wet-weighed (adjusting for water content) to avoid killing any microorganisms present while retaining as many fine particles as possible. The treatments were conducted using freshly collected sediments to be able to assess any physical (e.g., abrasion) and biological (e.g., pathogenic) stress responses.

A subset of the sediments was analyzed for grain size by sieving and pipetting after drying the sediments thoroughly in an oven. Grain size limits were defined according to the Udden-Wentworth US standard classification [50], while statistical parameters were determined using graphical measures from the cumulative size distributions as shown in S1 Fig. A summary of these parameters are provided in S1 Table. Reef sediments can be described as a well-sorted medium sand with virtually no silt or gravel. Conversely, Port sediments can be described as a poorly-sorted, medium sand with small amounts of gravel (approx. 1.97% >2000 μm) and silt and clay-sized particles (also referred as “mud”, approx. 1.25% <63 μm). No chemical or organic matter analysis of the sediments were conducted.

### Experimental setup

Fifty 8-day-old larvae were added to individual glass vials containing 45 mL filtered seawater (5 μm). Vials were allocated randomly to one of five different treatments: (1) a control treatment (no sediment), (2) a low “Reef” suspended sediment treatment (20 mg/L of homogenized sediment from Horseshoe Reef added), (3) a low “Port” sediment treatment (20 mg/L sediment from the Port site added), (4) a high “Reef” sediment treatment (100 mg/L sediment from Horseshoe reef added), and (5) a high “Port” sediment treatment (100 mg/L sediment from the Port site added). Nine to ten replicates were used per treatment. Vials were attached to a titer plate shaker set to 300 rpm to continuously maintain sediments in suspension for 24 h. Initial trials were conducted to assess the best exposure times that caused minimal mortality in control larvae and maintained both fine and coarse sediment particles in continuous suspension.

Treatments were chosen to characterize sub-lethal and lethal responses to suspended sediment concentrations for “tolerant” versus “sensitive” species as described by [12]. In addition, our goal was to test a low-concentration of suspended sediment (20 mg/L) representing typical conditions near dredging operations. Jones et al. [13] analyzed suspended sediment concentrations over a 30-day running mean period during the Barrow Island dredging project in Australia and found that the 95th percentile was 21 mg/L for 7 sites located at distances <1 km from the dredging. Similarly, although our 100 mg/L (high-dose) suspended sediment treatments were approximately 5 times higher than those typically observed during dredging operations, such conditions do occur over short periods (see [51]).

### 24 hour treatment exposure: larval survival assays

After 24 h of exposure, swimming larvae remaining in each vial were pipetted out, counted and examined with a microscope to determine the proportion that had survived. The vial was also carefully examined for potential settlers. A subset of larvae (N=6 per vial) was used to assess changes in respiration rates. Remaining larvae were allowed to recover for one week in filtered seawater (without sediments) in the same vials before being transported to the Florida Keys to conduct the settlement assays.

### 1-week post-exposure: larval settlement assays

Settlement assays were conducted by pooling all the larvae remaining in each treatment and then redistributing N=15 larvae to individual vials containing filtered seawater and a piece of rubble collected from Horseshoe reef to act as settlement cue. A total of 10 replicate glass vials per treatment were used to quantify larval settlement per treatment. After 24 h, the total number of larvae still swimming, or that had settled on the rubble, was manually scored using fluorescence microscopy. Larvae were classified as “settlers” only if they displayed visible signs of settlement (attachment to the substrate) and metamorphosis (i.e., transition from pear-shaped to flat/disc shape). To determine the cumulative percentage of larval settlement, the total number of settled corals (spat) was divided by the initial number of larvae added to each container.

### 24 hour and 1-week assessment of coral larval respiration

Respiration was measured using a microplate reader developed by Loligo Systems (Denmark), which allows measuring real-time respiration rates in each one of 24 wells using an optical fluorescence oxygen sensing technology (SDR SensorDish Reader®, PreSens, Germany) as described in [52–54]. This system comes pre-calibrated specifically for the reader used, and oxygen solubility is continuously calculated with oxygen sensors (optrodes) attached to the bottom of each well. Larval respiration rates were assessed at two time points, immediately after the 24 h treatment exposure and 1-week post-exposure. At each time point, larvae (N=6) were taken at random from each vial and placed in each of the wells of a sealed glass 125-µL microplate (without any sediments). A total of 8-10 replicate wells were used per treatment. Four additional control wells were used in each plate without any larvae to measure any background respiration in the treatment water.

Larvae were visually inspected for normal swimming behavior prior to each run. After each plate was prepared, wells were inspected to ensure there were no air bubbles, and an oxygen-impermeable seal was created using a silicone membrane covered with parafilm and a compression block. To maintain constant temperature, the microplate was placed inside a temperature-controlled flow-through water bath (included as part of the system), in a light-controlled room. The plate was then placed in a titer shaker at a low speed to maintain constant water motion. Finally, measurements of oxygen concentrations were taken every 15 seconds throughout the duration of the run and recorded using the SDR v 4.0.0 software (PreSens, Germany). Oxygen concentration was plotted as a function of time for individual wells and the first 10-20% (pO_2_ in % air saturation) linear decreases in oxygen were used to calculate the oxygen consumption rate (an example plot of oxygen consumption over time for 6 *O. faveolata* larvae placed in a single well of the microplate is shown in S2 Fig). Any portion of the slope that dipped below 70% air saturation was not used for analysis. Respiration rates were then corrected for background respiration and calculated as nanomoles of oxygen consumed per larva per minute.

### 16S sequencing of sediment samples

Immediately upon collection, ∼1 gram from each of the Port and Reef sediment samples collected per site was preserved in RNAlater for 16S sequencing. The DNA was extracted and PCR amplified with 16S rRNA gene V4 primers [55] with the Platinum Hot Start PCR Master Mix (2X) (ThermoFisher Scientific, Waltham, MA). The master mix was a 48-µl reaction and 2 µl of DNA template was used. The DNA was then amplified as follows: 94 °C for 3 minutes (1X), 94 °C for 45 seconds (35X), 50 °C for 60 seconds (35X), 72 °C for 90 seconds (35X), and 72 °C for 10 minutes (1X). Amplified products were cleaned using AMPure XP beads (Beckman Coulter, Brea, CA), normalized to 4 nM, and 5 µl of each normalized sample was pooled. The samples were sequenced on a MiSeq with PE-300v3 kits at the Halmos College of Natural Sciences and Oceanography at Nova Southeastern University.

### Statistical analysis

#### Larval performance

All analyses of larval performance data were done in R v.4.1.2. The effects of the sediment treatments on the proportion of larval survivorship and settlement were analyzed using binomial generalized linear models (GLM) with the lme4 package [56]. The models included sediment treatment as a fixed effect and replicate (vial) as a random effect. Model predictions (survivorship and settlement probabilities as well as the odds ratio between the control and the treatments) were plotted using the plot_model function from the sjPlot package [57] including 95% confidence intervals. Larval respiration rates after 24 h of treatment exposure and 1-week post-exposure were analyzed using a linear mixed-effects model (LMEM) with the lme4 package. The model included sediment treatment and time point (exposure and recovery) as fixed interactive effects, and vial, plate, and well as random effects. All data and code for larva survivorship, settlement and oxygen consumption can be found at Zenodo [58].

#### Microbiome analysis of sediment samples

DNA sequences were demultiplexed and further processed with Qiime2-2022.11 [59]. Since the reverse reads were of poor quality only the forward reads were analyzed. Using the program Cutadapt the primers were removed [60], and Amplicon Sequence Variants (ASVs) were generated with the program DADA2 [61] trimmed at the 10 and 205 bp positions. The ASVs were taxonomically classified with the classify-consensus-vsearch function and the silva 138 99 database [62]. If the sequences were annotated as chloroplast or mitochondria they were removed from the analysis. The data were then uploaded to R and converted into a Phyloseq v1.26.1 data object for analysis and graphical display of phylogenetic sequencing data.

Alpha-diversity was evaluated by rarefying to a minimum sequence depth of 60,000. Three alpha-diversity metrics, richness, Shannon-Wiener, and the Inverse Simpson index, were generated with the phyloseq estimate_richness function. The respective values were then tested for normality using qqnorm, qqline, and the Shapiro test. Upon passing normality, significance between sites was tested with an Analysis of Variance (ANOVA).

For beta-diversity, ASVs were filtered if they summed to <10 in 30% of the data. The filtered data were then transformed to centered log ratios (CLR) using the package Microbiome v0.9.99. The Vegan v2.5.4 package was used for beta-diversity testing for both within and between group differences. Dispersion of the samples (within group) was tested by using the function Vegdist (method = “Euclidean”, Permutations = 999) and Betadisper and was tested for significance with a permutation test. A permutational multivariate analysis of variance (PERMANOVA, function Adonis, method = “Euclidean”, Permutations = 999) was used to evaluate significance between groups.

To identify differentially abundant microbial ASVs between sites, the program Analysis of Compositions of Microbiomes with Bias Correction (ANCOM_BC) was used [63]. ANCOM_BC was used with an alpha <0.001 and a W statistic was >90. The ASVs were further evaluated if they had a log-fold change < −3.5 or > 3.5.

## Results

### Larval performance

Sediment treatments significantly impacted the performance of *O. faveolata* larvae (Fig 2, S2 Table). In general, sediment exposure negatively impacted larval survival and settlement for both Port and Reef sediment types. However, the effect of sediment concentration was important depending on the sediment type used in the treatment. Whereas both high and low Port sediment concentrations negatively impacted larval survival and settlement, only the high-dose Reef sediment treatment resulted in adverse effects (Figs 2 and S3). Specifically, survival proportions after 24 h in the treatments were above 93% for Control and Low Reef, but exposure to Low Port, High Reef, and High Port reduced survival probabilities to 90%, 86%, and 81%, respectively (p<0.05, df = 4, S2 and S3 Tables), resulting in significantly lower odds ratios of survivorship for these treatments (<0.6) compared to the Control (S3 Fig). Larval settlement probabilities 1-week after treatments were 21-22% in the Control and Low Reef treatments, but were reduced to less than 12% in the Low Port, High Reef, and High Port treatments (p<0.05, df = 4, S2 and S3 Tables). This resulted in lower (<0.5) odds ratios for settlement in the Low Port, High Reef, and High Port treatments compared to the Control (S3 Fig).

**Fig 2.**
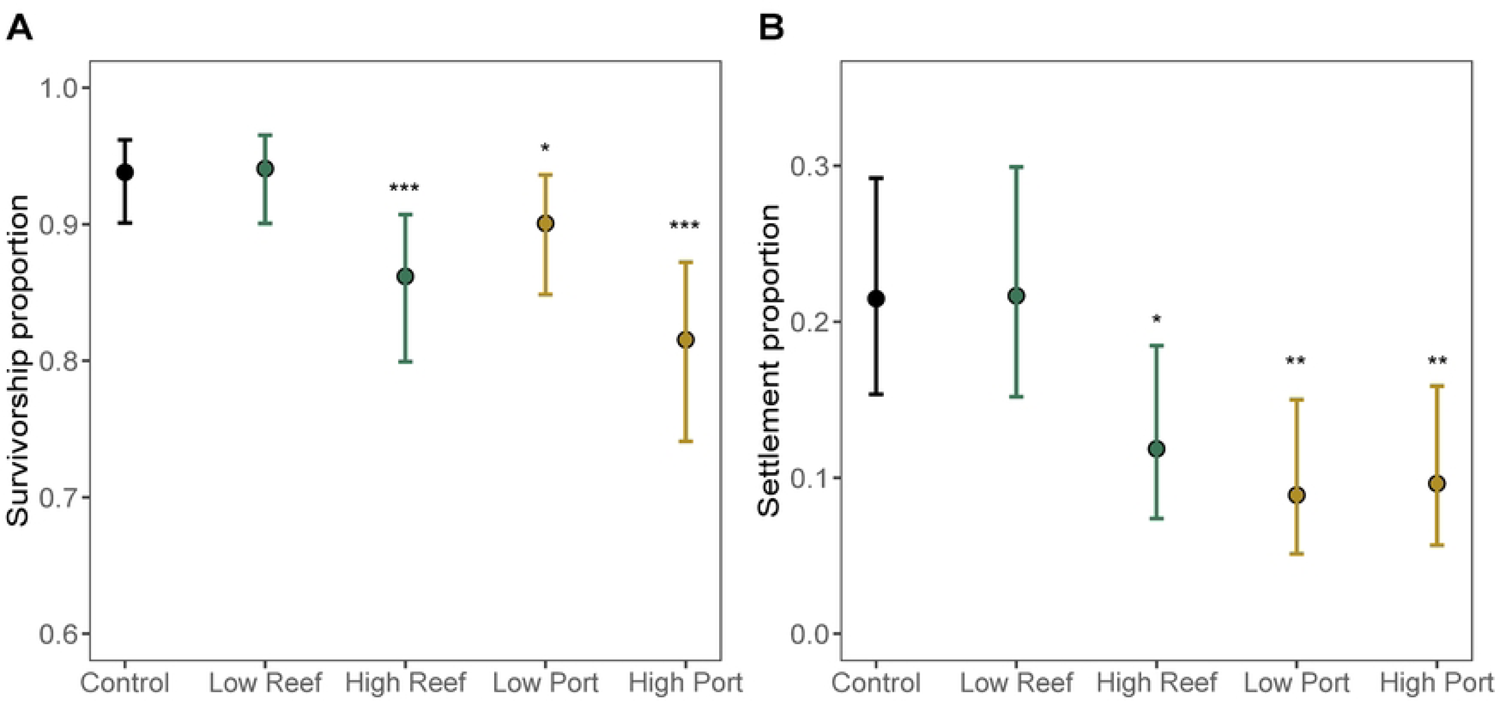
Effects of sediments on *Orbicella faveolata* larvae (model estimated values ± 95% CI). (A) Proportion of larval survival after a 24 h exposure to experimental treatments. (B) Proportion of larval settlement after one week of recovery from experimental treatments. Asterisks denote significant differences in a sediment treatment compared to controls with no sediment added (*: p < 0.05; **: p < 0.01; ***: p < 0.001). Colors match Figs 3 and 4 and denote either Port or Reef sediment treatments.

None of the sediment treatments affected the larval respiration rates at either of the time points (S4 Table, S4 Fig). Average respiration rates per larva were 9.33 x 10^-4^ ± 2.53 x 10^-4^ nmol O_2_ min^-1^ (mean ± SD) across treatments after 24 h of exposure and declined to 6.01 x 10^-4^ ± 1.97 x 10^-4^ nmol O_2_ min^-1^ after a week of recovery, but this was only marginally significant (p<0.06, df = 1, F = 15.83, S4 Table).

### Microbiome analysis

A total of seven sediment samples (Port N=3 and Reef N=4) resulted in sequence counts between 93,626 and 419,403 (median=171,281). After processing and filtering, 1,852 ASVs were used for the analysis. The median alpha-diversity was slightly higher in the Reef sediments (richness: Reef=3604 vs Port=3001, Shannon-Wiener: Reef=7.26 vs Port=7.20, and Inverse Simpson: Reef=654.24 vs Port=584.8), but this difference was not significant between sites (Fig 3A). In contrast, within-group (permutest, p=0.023) and between-group beta-diversity were significantly different (PERMANOVA, Fmodel=6.127, R^2^=0.55 p=0.03, Fig 3B), with the microbial communities in Reef sediments having a more dispersed beta-diversity compared to Port sediments (average distance to centroid: Reef=55.76 vs Port=41.99). Of the seven most relatively abundant bacteria orders, the Reef and Port sediments shared four taxa (57%) with NB1-J and Cyanobacteriales being unique to Reef sediments, and Desulfobacterales being unique to the Port sediments (Fig 3C).

**Fig 3.**
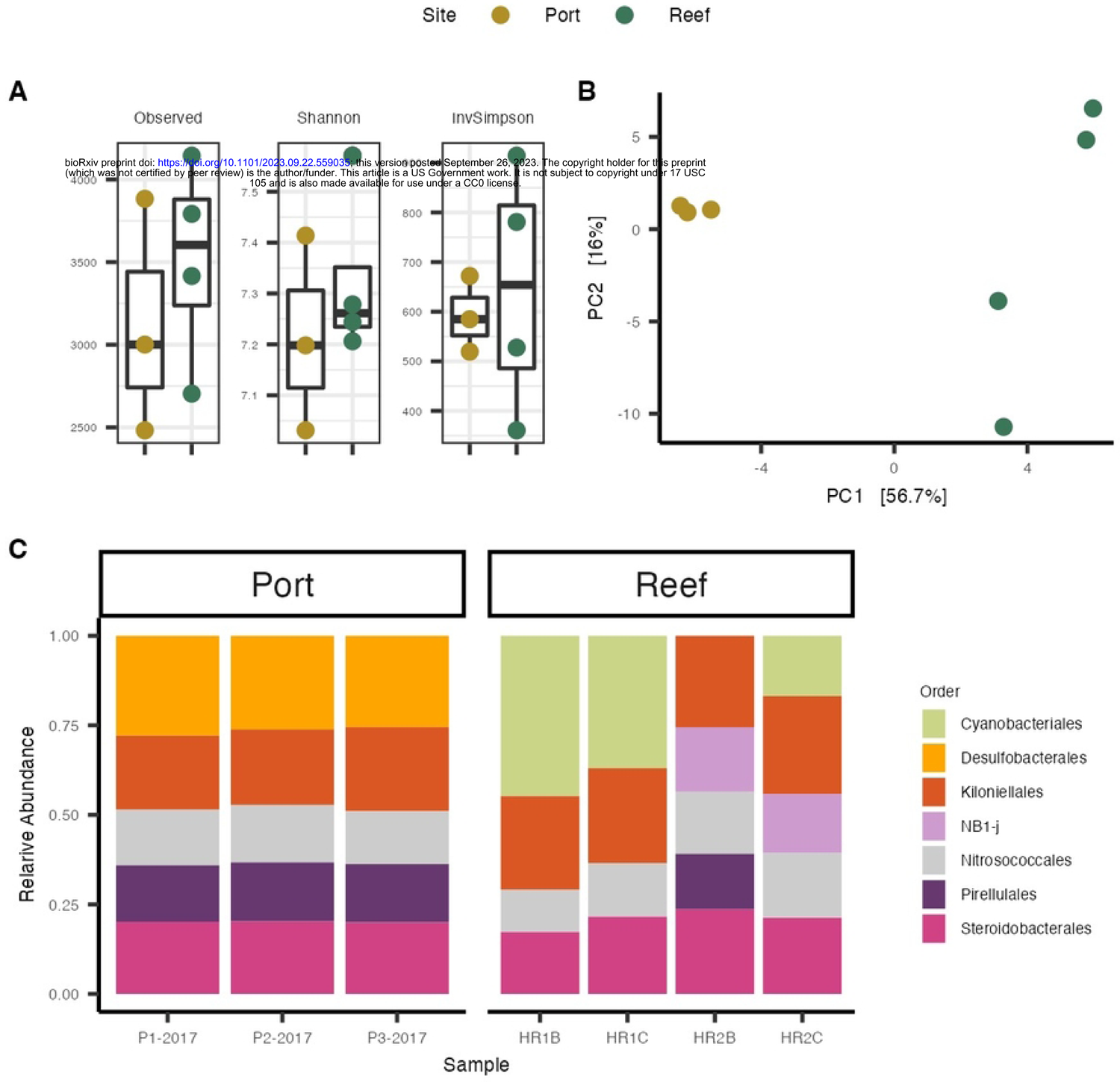
Microbial communities found in Port and Reef sediments. (A) Alpha diversity differences between Reef and Port samples using three diversity metrics: (1) Species richness (2) Shannon diversity Index, and (3) Inverse Simpson Index. (B) Beta diversity of microbial composition between Port and Reef sediment samples. (C) The cumulative relative abundances of the most abundant (0.05>) microbial order per sample grouped by the site of sediment collections.

Differential abundance analysis was conducted to identify the differences between sites, resulting in 293 significant ASVs (Fig 4). Of these, there were a total of 74 orders, 82 families, and 96 genera. The most prevalent differentially abundant orders were Steroidobacterales (N=29), Desulfobacterales (N=19), and Pirellulales (N=15). The Port sediments had more enriched ASVs (N=223) compared to Reef sediments (N=70). More specifically, the ASVs most enriched in Port sediments were from the orders Ectothiorhodospirales (family *Ectothiorhodospiraceae*, genus *Thiogranum*), an uncultured genus from the Class Gammaproteobacteria, Desulfobacterales (*Desulfosarcinaceae*, Sva0081 sediment group), and Steroidobacterales (*Woeseiaceae*, *Woeseia*). In the Reef sediments, the most enriched orders were Alteromonadales (*Moritellaceae* from an uncultured genus), Steroidobacterales *(Woeseiaceae*, *Woeseia*), and Nitrosococcales (*Nitrosococcaceae*, *AqS1*).

**Fig 4.**
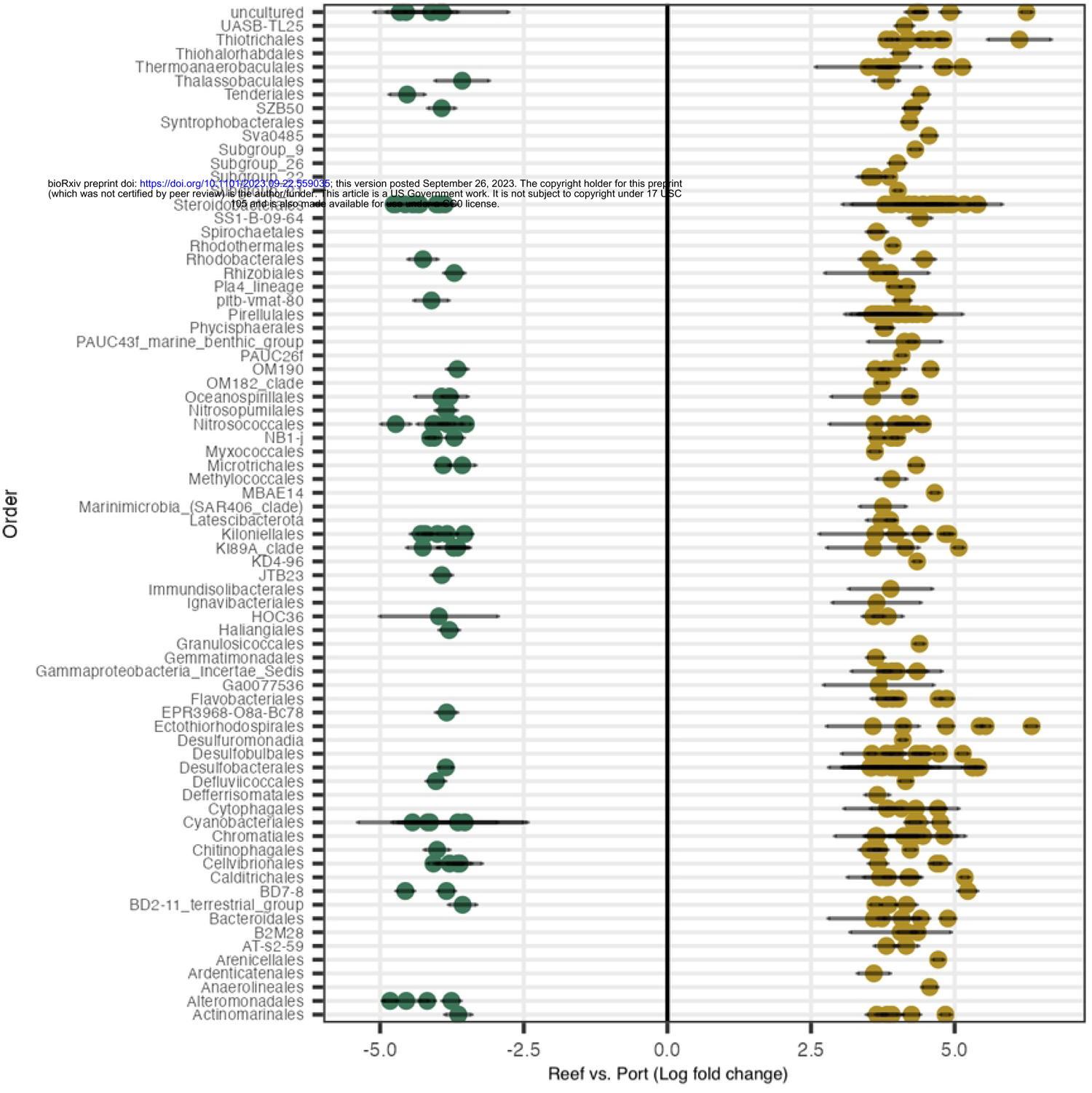
Microbial Amplicon Sequence Variants (ASVs) associated with Reef or Port sediments. Differential abundances between Reef and Port, where the y-axis shows significant ASVs (padj<0.001, W statistic >90) grouped by order. Only ASVs with a log-fold change <-3.5 and >3.5 were visualized.

## Discussion

In this study, we investigated differences in the survival and settlement of larvae from the threatened Caribbean coral *Orbicella faveolata* after short-term exposure to sediments collected near the recently dredged Port Miami channel (considered anthropogenically modified) versus sediments collected from the natal reef of the parent colonies in the Florida Keys (considered natural or unmodified). Neither of the sediment types were dried or sterilized to avoid changing their biological or chemical properties. Overall, we found that although *O. faveolata* larvae exposed to high doses of both sediment types (“Port” and “Reef”) exhibited adverse survival and settlement rates, only the Port sediments resulted in adverse effects when exposed to low-dose treatments, characteristic of conditions near actual dredging projects (e.g., [13]). The low-dose Reef sediment treatment was not detrimental to coral larvae, and showed similar outcome odds of survival and settlement compared to the no sediment (control) treatment (see S3 Fig).

Port sediments had different microbiome profiles and significantly higher abundances of potentially pathogenic bacteria compared to Reef sediments, which may explain the reduced larval survival and settlement observed even in the low-dose treatment. We speculate that Port sediments could have been more detrimental to larvae compared to reef sediments due to differences in their physical and chemical composition (e.g., grain size and sediment composition), and other biological differences (e.g., different microbial communities). Together, these findings suggest that, although suspended sediments may be generally detrimental to the survival and recruitment potential of *O. faveolata* larvae, anthropogenic sediments, such as those occurring adjacent to recent dredging projects, may incur greater harm compared to natural reef sediments.

### Effects on larval survival

Both of our sediment sources significantly impacted the survival of *O. faveolata* larvae, however, exposure to both the low- and high-dose Port sediment treatments reduced survival proportions to 86% and 81%, respectively (compared to the control treatment), while the high-dose Reef sediment treatment only reduced the survival proportion to 90% (Fig 2A). Suspended sediments may reduce larval survival through physical abrasion [38] and reduced light over prolonged periods can also lead to decreased photosynthetic efficiency of larval symbionts and subsequent larval mortality from starvation [23]. However, in our study, decreased light was unlikely the driver of our results, given the short exposure to sediments (24 h) and the use of aposymbiotic (symbiont-free) larvae. Instead, we suspect that reduced larval survival in both of our high-dose sediment treatments (Fig 2A) could have been caused by continuous physical abrasion from suspended sediments during exposure. Given the pelagic nature of coral larvae, this stage is likely to be more prone to physical abrasion from moving sediment particles compared to coral recruits, on which sediment can accumulate on the growing coral tissue.

Increased mortality of *O. faveolata* larvae in both of our Port sediment treatments (compared to Reef sediments) could have also been due to chemical or microbial differences in the sediment composition and/or grain size. A recent study [32] assessed the effects of sedimentation on recruits from the brooding coral *Porites astreoides* using either reef sediments collected from the site of adult colonies, or sediments collected near Port Everglades (Fort Lauderdale, Florida) to resemble the grain size composition often found in dredging areas. Similar to our study, the authors found that reef sediments did not negatively affect coral survival, but Port sediments did. However, since there were only trace amounts of mud-sized particles in our Port sediments (approximately 1.25%), we hypothesize that Port sediments may be more likely to have affected coral larvae through microbial processes leading to decreased survival.

### Effects on larval settlement

Except for the low-dose Reef sediment treatment, we showed significant effects of both sediment types (Port and Reef) after one week of sediment exposure, resulting in approximately 50-60% reduction in settlement in *O. faveolata* larvae compared to the control treatment (Figs 2B and S1). Reduced larval settlement has been documented for a few Pacific coral species in the presence of suspended sediments (in concentrations less than 10 mg/L, [23]), presumably due to changes in light quality and quantity (as a result of increased light attenuation). Sediment-covered surfaces have also been shown to decrease larval settlement in various coral species such as *Pocillopora damicornis* [64], *Acropora digitifera* [38], *Acropora millepora* [38], and *O. faveolata* [31], likely due to limited space to settle (preventing larvae from accessing and attaching to suitable substrate), higher risk of abrasion, smothering/burial by deposited sediments, and masking of appropriate chemical cues for settlement [23].

An important distinction of our study compared to previous work on the effects of sedimentation on coral larvae is that we tested potential latent effects of sediment exposure on eventual settlement one week after acute exposure. Thus, the settlement environment (i.e., light and condition or exposure of settlement substrates) were the same across treatments, and consequently neither sediment deposition on the coral surfaces or settlement substrates, nor changes in light, were drivers of the reduced settlement observed in the exposed larvae. Furthermore, since our low-dose Reef sediment treatment was not detrimental to coral settlement (but the low-dose Port sediment treatment was), we speculate that disruption of potential settlement cues in larvae exposed to Port sediments may have resulted from biological/chemical damage (potential interactions with any contaminants or bacterial pathogens present in the Port sediments). Sediment-stressed larvae could also have had lower energy reserves available for allocation to settlement and subsequent metamorphosis, although no significant respiration rate differences were observed among treatments (see S4 Fig). One reason why our study may have found latent effects is that previous studies conducted experiments using sediments that had been dried prior to exposure. Oven drying can change the chemical composition of the sediments and kill bacteria [5], masking these potentially important biological or chemical effects. The inclusion of wet sediments in our study may therefore have helped preserve these properties and allowed us to explore how bacterial assemblages can affect larval survival and settlement.

### Microbiome analysis of sediment samples and potential implications for coral survival and settlement

Coral settlement is the result of complex behavioral responses that are modulated by distinct microbial signatures in CCA [40–43]. In this study, we hypothesized that microbial differences among the two sediment types may have resulted in decreases in survivorship and potential impairment of the ability of larvae to detect cues for settlement and metamorphosis. The microbiome analysis of the sediment samples collected near the Port revealed differences in microbial beta-diversity and differentially abundant bacteria taxa compared to Reef sediments (Figs 3BC and 4). Relative abundance analysis of the most abundant taxa showed that Desulfobacterales were unique to the Port sediments (Fig 3C). These taxa have been found to be enriched in microbial communities from other inlets in southeast Florida [65] and have also been associated with black band disease [66], suggesting that this bacterial taxa may play a role in coral health and its enrichment in Port sediments may have contributed to the reduced survival and settlement in larvae exposed to both low- and high-doses of these sediments (Fig 2A). Moreover, Desulfobacterales are anaerobic sulfate-reducing bacteria [67] that reduce sulfate (SO_4_^2−^) to hydrogen sulfide (H_2_S). Desulfobacterales are abundant in organic-rich and high nutrient environments linked to eutrophication [68], and their waste product, H_2_S, is correlated with coral tissue degradation [11], and algal mortality [69]. Port sediments enriched in Desulfobacterales (Fig 4) may therefore reflect elevated H_2_S and stressful hypoxic conditions that may have resulted in lower survival and settlement rates.

Sediments may also act as vectors for the transmission of diseases such as Stony Coral Tissue Loss Disease [22], which is documented to have appeared for the first time near Port Miami in 2014 [70, 71]. Also, bacteria genera from two common human pathogens, *Staphylococcus* and *Streptococcus*, have been found across multiple time points in surface water and sediment samples near Port Everglades [65, 72], another southeast Florida inlet subject to dredging impacts. Our findings add to these studies and also suggest that port inlets may carry potentially pathogenic bacteria which may be transported to nearby reefs through various sediment transport processes (natural or anthropogenic), resulting in potential exposure to coral larvae and other downstream health impacts.

### Effects on coral larval respiration

Despite significant effects of sedimentation on *O. faveolata* larval survival and settlement (Figs 2 and S3), none of the sediment treatments had significant effects on coral larval respiration (S4 Fig and S4 Table). Sedimentation has been shown to decrease photosynthesis to respiration (P/R) ratios in adult corals, via decreased photosynthesis of the algal symbionts and increased respiration of the host (e.g., [73]). However, in this study, we used aposymbiotic larvae, which may explain the lack of significant effects on respiration. Alternatively, changes in respiration may have not been detected because it is not possible to maintain the sediment treatments in the microplate reader while respiration measurements are taken. Lower respiration rates across all treatments at the 1-week recovery period (compared to the 24-hr sediment exposure, see S3 Fig) could have resulted from ontogenetic and/or behavioral differences during those two time points (active swimming versus exploration of suitable substrate for settlement), e.g. [74]).

### Management implications for the conservation of corals

Our study is the first to examine the effects of suspended sediments on larvae from an Atlantic coral species. With the two major commercial shipping ports in southeast Florida planning major expansions which require multi-year dredge projects, this information is important for understanding the impacts of sedimentation on susceptible life stages of corals, and assisting with the development of protective management strategies. *O. faveolata* is listed under the US Endangered Species Act (ESA) and already shows poor recruitment [49, 75, 76]. Recent work confirms the ESA Critical Habitat (legal) requirements that larvae require substrates free of sediment for settlement [31]. Importantly, our study showed that while sediments have detrimental effects on the survival and settlement of *O. faveolata* larvae at high doses, even brief exposures to low doses of anthropogenic sediments found near a recently dredged Port during the pelagic phase could reduce larval survival and disrupt settlement, potentially compromising the recruitment success and recovery potential of this species. Furthermore, we showed substantial direct mortality from short-term sediment exposure, but also an additional significant latent effect on the surviving larvae’s capacity to successfully settle one week after sediment exposure. These results indicate that sediments which remain adjacent to a dredging project even after ∼2 years may still have significant effects on early life stages of corals during relatively short-term resuspension events (e.g., storms). This highlights the potential for long-term impacts of dredging (e.g., critical habitat modification) as a result of resuspension of anthropogenic sediments. However, it is unclear how long such sediments can persist in the area, or how often resuspension events might happen.

Our findings also highlight the importance of setting temporary moratoriums as a management tool during dredging operations. Since 1993 dredging projects in Western Australia conducted near coral reefs have been required to implement a coral environmental (“stand-down”) window spanning five days before spawning to seven days after spawning [13, 77]. The *O. faveolata* larvae used in our experiments were eight days old and were adversely impacted by both sediment types with just 24 h of exposure, with detrimental latent effects still detectable after one week. Consequently, a one-week “stand down” period may be insufficient for broadcast spawner coral species such as *O. faveolata* given the length of time it takes for larvae to become competent and find suitable substrate for settlement. Based on our results, we recommend setting environmental windows which encompass the months of peak spawning (typically August and September for *O. faveolata* [78, 79]), plus an additional period of at least two to three weeks after spawning to prevent negative effects on settlement. These months are also typically the peak period for heat-stressed induced coral bleaching, and avoiding dredging during these months would therefore have multiple benefits.

Finally, we showed that sediments collected near the Port contained higher abundances of potentially pathogenic bacteria compared to Reef sediments, aligning with previous work conducted in the vicinity of the Port Everglades inlet [65, 72]. These studies highlight the importance of monitoring for potential contaminants in sediments and changes in the microbiome which may result from dredging. In addition, although we only looked at the effects of suspended sediments on coral larval performance, suspended sediments can act in concert with other stressors typically associated with dredging. For example, [51] showed that elevated suspended sediment concentrations, when simultaneously combined with reduced light levels, resulted in partial mortality for multiple Pacific coral species. These findings emphasize the importance of monitoring both suspended sediments and benthic light availability (PAR) during large-scale dredging projects, and highlight the need for similar studies for Atlantic coral species.

## Acknowledgements

The authors thank C. Sinigalliano, J. Hendee, B. Jensen, J. Destache, C. Pasparakis, and B. Young for help in the laboratory. Field assistance was provided by scientists from the NOAA Southeast Fisheries Science Center (D. Williams, A. Bright and A. Peterson). J. Wolfe provided GIS assistance with Fig 1. *O. faveolata* gametes were collected under permit FKNMS-2016-047-A1 to M.W. Miller. Sediment collections were determined to be exempt from requirements for an Environmental Resource Permit (ERP) by the Florida’s Department of Environmental Protection (DeMinimus exemption). This research was carried out in part under the auspices of the Cooperative Institute for Marine and Atmospheric Studies (CIMAS), a cooperative institute of the University of Miami and the National Oceanic and Atmospheric Administration (NOAA), Cooperative Agreement NA 20OAR4320472. The scientific results and conclusions, as well as any opinions expressed herein, are those of the author(s) and do not necessarily reflect the views of NOAA or the Department of Commerce.

## Supporting information

**S1 Fig. Composite display of grain size analysis of sediment samples.** Two replicates per sediment type (Port versus Reef) were used.

**S2 Fig. Example plot of oxygen consumption over time for 6 *Orbicella faveolata* larvae placed in a single well of the microplate.** The R^2^ value is given for the linear trend. The first 10–20% (pO2 in % air saturation) linear decreases in oxygen were used to calculate respiration rates per larva. Any portion of the slope that dipped below 70% air saturation was not used for data analysis.

**S3 Fig. Odds ratios (OR) for the association between *Orbicella faveolata* larvae exposure to sediment treatments.** (A) larval survival after 24 h of treatments, and (B) larval settlement after one week of recovery from experimental treatments. OR=1 Exposure does not affect odds of outcome, OR>1 Exposure associated with higher odds of outcome, and OR<1 Exposure associated with lower odds of outcome.

**S4 Fig. Respiration rates (mean ± 95 CI) in *Orbicella faveolata* larvae** after a 24 h exposure to experimental treatments (left panel) and after one week of recovery from experimental treatments (right panel). White dots denote individual replicate wells.

**S1 Table. Size fraction analysis of sediment samples.** Values correspond to graphic in S1 Fig.

**S2 Table.** Generalized linear mixed-effect models outputs for *Orbicella faveolata* larvae performance. DF: Degrees of freedom

**S3 Table. Results of the final generalized linear mixed-effect models for *Orbicella faveolata* performance.** A) larval survival after 24 h of treatments, and B) larval settlement after one week of recovery from experimental treatments (shown in Fig 1). The coefficient estimates describe the change in the log odds for each treatment level compared to the base level (control). Asterisks denote significant differences in a sediment treatment compared to controls with no sediment added (*: p < 0.05; **: p < 0.01; ***: p < 0.001).

**S4 Table.** Linear mixed-effect model outputs for *Orbicella faveolata* larvae respiration.

## References

1. Cunning R, Silverstein RN, Barnes BB, Baker AC. Extensive coral mortality and critical habitat loss following dredging and their association with remotely-sensed sediment plumes. Mar Pollut Bull. 2019;145:185–99. Epub 2019/10/09. doi: 10.1016/j.marpolbul.2019.05.027. PubMed PMID: 31590775.

2. Foster T, Corcoran E, Erftemeijer P, Fletcher C, Peirs K, Dolmans C, et al. Dredging and port construction around coral reefs. PIANC Environmental Commission. Report, 2010.

3. Miller MW, Karazsia J, Groves CE, Griffin S, Moore T, Wilber P, et al. Detecting sedimentation impacts to coral reefs resulting from dredging the Port of Miami, Florida USA. PeerJ. 2016;4:e2711. Epub 2016/11/30. doi: 10.7717/peerj.2711. PubMed PMID: 27896036; PubMed Central PMCID: PMCPMC5119242.

4. Rosales SM, Sinigalliano C, Gidley M, Jones PR, Gramer LJ. Oceanographic habitat and the coral microbiomes of urban-impacted reefs. PeerJ. 2019;7:e7552. Epub 2019/10/01. doi: 10.7717/peerj.7552. PubMed PMID: 31565557; PubMed Central PMCID: PMCPMC6743471.

5. Rushmore ME, Ross C, Fogarty ND. Physiological responses to short-term sediment exposure in adults of the Caribbean coral Montastraea cavernosa and adults and recruits of Porites astreoides. Coral Reefs. 2021;40(5):1579–91. doi: 10.1007/s00338-021-02156-0.

6. Walker BK, Gilliam DS, Dodge RE, Walczak J. Dredging and shipping impacts on southeast Florida coral reefs. Proceedings of the 12th International Coral Reef Symposium, Cairns, Australia, 9-13 July 2012. 2012.

7. Jones R, Bessell-Browne P, Fisher R, Klonowski W, Slivkoff M. Assessing the impacts of sediments from dredging on corals. Mar Pollut Bull. 2016;102(1):9–29. Epub 2015/12/15. doi: 10.1016/j.marpolbul.2015.10.049. PubMed PMID: 26654296.

8. Weber M, Lott C, Fabricius KE. Sedimentation stress in a scleractinian coral exposed to terrestrial and marine sediments with contrasting physical, organic and geochemical properties. Journal of Experimental Marine Biology and Ecology. 2006;336(1):18–32. doi: 10.1016/j.jembe.2006.04.007.

9. Jones R, Fisher R, Bessell-Browne P. Sediment deposition and coral smothering. PLoS One. 2019;14(6):e0216248. Epub 2019/06/20. doi: 10.1371/journal.pone.0216248. PubMed PMID: 31216275; PubMed Central PMCID: PMCPMC6584000

10. Storlazzi CD, Norris BK, Rosenberger KJ. The influence of grain size, grain color, and suspended-sediment concentration on light attenuation: Why fine-grained terrestrial sediment is bad for coral reef ecosystems. Coral Reefs. 2015;34(3):967–75. doi: 10.1007/s00338-015-1268-0.

11. Weber M, de Beer D, Lott C, Polerecky L, Kohls K, Abed RM, et al. Mechanisms of damage to corals exposed to sedimentation. Proc Natl Acad Sci U S A. 2012;109(24):E1558–67. Epub 2012/05/23. doi: 10.1073/pnas.1100715109. PubMed PMID: 22615403; PubMed Central PMCID: PMCPMC3386076.

12. Erftemeijer PL, Riegl B, Hoeksema BW, Todd PA. Environmental impacts of dredging and other sediment disturbances on corals: a review. Mar Pollut Bull. 2012;64(9):1737–65. Epub 2012/06/12. doi: 10.1016/j.marpolbul.2012.05.008. PubMed PMID: 22682583.

13. Jones R, Ricardo GF, Negri AP. Effects of sediments on the reproductive cycle of corals. Mar Pollut Bull. 2015;100(1):13–33. Epub 2015/09/20. doi: 10.1016/j.marpolbul.2015.08.021. PubMed PMID: 26384866.

14. Piniak GA. Effects of two sediment types on the fluorescence yield of two Hawaiian scleractinian corals. Mar Environ Res. 2007;64(4):456–68. Epub 2007/06/15. doi: 10.1016/j.marenvres.2007.04.001. PubMed PMID: 17568664.

15. Barnes BB, Hu C, Kovach C, Silverstein RN. Sediment plumes induced by the Port of Miami dredging: Analysis and interpretation using Landsat and MODIS data. Remote Sensing of Environment. 2015;170:328–39. doi: 10.1016/j.rse.2015.09.023.

16. Fisher R, Stark C, Ridd P, Jones R. Spatial Patterns in Water Quality Changes during Dredging in Tropical Environments. PLoS One. 2015;10(12):e0143309. Epub 2015/12/03. doi: 10.1371/journal.pone.0143309. PubMed PMID: 26630575; PubMed Central PMCID: PMCPMC4667927

17. Eggleton J, Thomas KV. A review of factors affecting the release and bioavailability of contaminants during sediment disturbance events. Environ Int. 2004;30(7):973–80. Epub 2004/06/16. doi: 10.1016/j.envint.2004.03.001. PubMed PMID: 15196845.

18. Jones RJ. Spatial patterns of chemical contamination (metals, PAHs, PCBs, PCDDs/PCDFS) in sediments of a non-industrialized but densely populated coral atoll/small island state (Bermuda). Mar Pollut Bull. 2011;62(6):1362–76. Epub 2011/05/10. doi: 10.1016/j.marpolbul.2011.01.021. PubMed PMID: 21549399.

19. Su SH, Pearlman LC, Rothrock JA, Iannuzzi TJ, Finley BL. Potential long-term ecological impacts caused by disturbance of contaminated sediments:a case study. Environ Manage. 2002;29(2):234–49. Epub 2002/01/30. doi: 10.1007/s00267-001-0005-3. PubMed PMID: 11815826.

20. Hodgson G. Tetracycline reduces sedimentation damage to corals. Marine Biology. 1990;104:493–6.

21. Voss JD, Richardson LL. Coral diseases near Lee Stocking Island, Bahamas: patterns and potential drivers. Diseases of Aquatic Organisms. 2006;69(1):33–40.

22. Studivan MS, Rossin AM, Rubin E, Soderberg N, Holstein DM, Enochs IC. Reef Sediments Can Act As a Stony Coral Tissue Loss Disease Vector. Frontiers in Marine Science. 2022;8. doi: 10.3389/fmars.2021.815698.

23. Tuttle LJ, Donahue MJ. Effects of sediment exposure on corals: a systematic review of experimental studies. Environ Evid. 2022;11(1):4. Epub 2022/02/15. doi: 10.1186/s13750-022-00256-0. PubMed PMID: 35154667; PubMed Central PMCID: PMCPMC8818373.

24. Ricardo GF, Jones RJ, Negri AP, Stocker R. That sinking feeling: Suspended sediments can prevent the ascent of coral egg bundles. Sci Rep. 2016;6:21567. Epub 2016/02/24. doi: 10.1038/srep21567. PubMed PMID: 26898352; PubMed Central PMCID: PMCPMC4761919.

25. Ricardo GF, Jones RJ, Clode PL, Humanes A, Negri AP. Suspended sediments limit coral sperm availability. Sci Rep. 2015;5:18084. Epub 2015/12/15. doi: 10.1038/srep18084. PubMed PMID: 26659008; PubMed Central PMCID: PMCPMC4677285.

26. Ricardo GF, Jones RJ, Clode PL, Humanes A, Giofre N, Negri AP. Sediment characteristics influence the fertilisation success of the corals Acropora tenuis and Acropora millepora. Mar Pollut Bull. 2018;135:941–53. Epub 2018/10/12. doi: 10.1016/j.marpolbul.2018.08.001. PubMed PMID: 30301119.

27. Babcock R, Davies P. Effects of sedimentation on settlement of Acropora millepora. Coral reefs. 1991;9:205–8.

28. Babcock R, Smith L, editors. Effects of sedimentation on coral settlement and survivorship. Proceedings of the Ninth International Coral Reef Symposium, Bali, 23-27 October 2000; 2000: Citeseer.

29. Perez K, 3rd, Rodgers KS, Jokiel PL, Lager CV, Lager DJ. Effects of terrigenous sediment on settlement and survival of the reef coral Pocillopora damicornis. PeerJ. 2014;2:e387. Epub 2014/06/03. doi: 10.7717/peerj.387. PubMed PMID: 24883248; PubMed Central PMCID: PMCPMC4034646.

30. Ricardo GF, Jones RJ, Nordborg M, Negri AP. Settlement patterns of the coral Acropora millepora on sediment-laden surfaces. Sci Total Environ. 2017;609:277–88. Epub 2017/07/28. doi: 10.1016/j.scitotenv.2017.07.153. PubMed PMID: 28750231.

31. Speare KE, Duran A, Miller MW, Burkepile DE. Sediment associated with algal turfs inhibits the settlement of two endangered coral species. Mar Pollut Bull. 2019;144:189–95. Epub 2019/06/11. doi: 10.1016/j.marpolbul.2019.04.066. PubMed PMID: 31179987.

32. Fourney F, Figueiredo J. Additive negative effects of anthropogenic sedimentation and warming on the survival of coral recruits. Sci Rep. 2017;7(1):12380. Epub 2017/09/30. doi: 10.1038/s41598-017-12607-w. PubMed PMID: 28959051; PubMed Central PMCID: PMCPMC5620051.

33. Moeller M, Nietzer S, Schils T, Schupp PJ. Low sediment loads affect survival of coral recruits: the first weeks are crucial. Coral Reefs. 2016;36(1):39–49. doi: 10.1007/s00338-016-1513-1.

34. Fabricius KE. Effects of terrestrial runoff on the ecology of corals and coral reefs: review and synthesis. Mar Pollut Bull. 2005;50(2):125–46. Epub 2005/03/02. doi: 10.1016/j.marpolbul.2004.11.028. PubMed PMID: 15737355.

35. Price N. Habitat selection, facilitation, and biotic settlement cues affect distribution and performance of coral recruits in French Polynesia. Oecologia. 2010;163(3):747–58. Epub 2010/02/20. doi: 10.1007/s00442-010-1578-4. PubMed PMID: 20169452; PubMed Central PMCID: PMCPMC2886133.

36. Ritson-Williams R, Paul VJ, Arnold SN, Steneck RS. Larval settlement preferences and post-settlement survival of the threatened Caribbean corals Acropora palmata and A. cervicornis. Coral Reefs. 2009;29(1):71–81. doi: 10.1007/s00338-009-0555-z.

37. Birrell CL, McCook LJ, Willis BL. Effects of algal turfs and sediment on coral settlement. Mar Pollut Bull. 2005;51(1-4):408–14. Epub 2005/03/11. doi: 10.1016/j.marpolbul.2004.10.022. PubMed PMID: 15757739.

38. Gilmour J. Experimental investigation into the effects of suspended sediment on fertilisation, larval survival and settlement in a scleractinian coral. Marine Biology. 1999;135:451–62.

39. Goh B, Lee C, editors. A study of the effect of sediment accumulation on the settlement of coral larvae using conditioned tiles. Proceedings Eleventh International Coral Reef Symposium, Ft Lauderdale, Florida; 2008.

40. Jorissen H, Galand PE, Bonnard I, Meiling S, Raviglione D, Meistertzheim AL, et al. Coral larval settlement preferences linked to crustose coralline algae with distinct chemical and microbial signatures. Sci Rep. 2021;11(1):14610. Epub 2021/07/18. doi: 10.1038/s41598-021-94096-6. PubMed PMID: 34272460; PubMed Central PMCID: PMCPMC8285400.

41. Quinlan ZA, Ritson-Williams R, Carroll BJ, Carlson CA, Nelson CE. Species-Specific Differences in the Microbiomes and Organic Exudates of Crustose Coralline Algae Influence Bacterioplankton Communities. Front Microbiol. 2019;10:2397. Epub 2019/11/30. doi: 10.3389/fmicb.2019.02397. PubMed PMID: 31781048; PubMed Central PMCID: PMCPMC6857149.

42. Siboni N, Abrego D, Puill-Stephan E, King WL, Bourne DG, Raina J-B, et al. Crustose coralline algae that promote coral larval settlement harbor distinct surface bacterial communities. Coral Reefs. 2020;39(6):1703–13. doi: 10.1007/s00338-020-01997-5.

43. Sneed JM, Ritson-Williams R, Paul VJ. Crustose coralline algal species host distinct bacterial assemblages on their surfaces. ISME J. 2015;9(11):2527–36. Epub 2015/04/29. doi: 10.1038/ismej.2015.67. PubMed PMID: 25918832; PubMed Central PMCID: PMCPMC4611515.

44. NOAA CD. Endangered and threatened wildlife and plants: Final listing determinations on proposal to list 66 reef-building coral species and to reclassify elkhorn and staghorn corals. Federal Register. 2014:53851–4123.

45. Vazquez C. Explosive report finds PortMiami dredging caused extensive coral reef damage. Local 10 News [Internet]. 2023 Sept 06. Available from: https://www.local10.com/news/local/2023/09/06/explosive-report-finds-portmiami-dredging-caused-extensive-coral-reef-damage/#:|:text=MIAMI%20%E2%80%93%20A%20new%20federal%20report,along%20the%20port's%20entrance%20channel.

46. Gintert BE, Precht WF, Fura R, Rogers K, Rice M, Precht LL, et al. Regional coral disease outbreak overwhelms impacts from a local dredge project. Environ Monit Assess. 2019;191(10):630. Epub 2019/09/15. doi: 10.1007/s10661-019-7767-7. PubMed PMID: 31520148.

47. May LA, Miller CV, Moffitt ZJ, Balthis L, Karazsia J, Wilber P, et al. Acute turbidity exposures with Port of Miami sediments impact Orbicella faveolata tissue regeneration. Marine Pollution Bulletin. 2023;193:115217.

48. Szmant AM, Miller MW, editors. Settlement preferences and post-settlement mortality of laboratory cultured and settled larvae of the Caribbean hermatypic corals Montastraea faveolata and Acropora palmata in the Florida Keys, USA. Proc 10th Int Coral Reef Symp; 2006.

49. Miller MW. Post-settlement survivorship in two Caribbean broadcasting corals. Coral Reefs. 2014;33(4):1041–6. doi: 10.1007/s00338-014-1177-7.

50. Wentworth CK. A scale of grade and class terms for clastic sediments. The journal of geology. 1922;30(5):377–92.

51. Bessell-Browne P, Negri AP, Fisher R, Clode PL, Duckworth A, Jones R. Impacts of turbidity on corals: The relative importance of light limitation and suspended sediments. Mar Pollut Bull. 2017;117(1-2):161–70. Epub 2017/02/07. doi: 10.1016/j.marpolbul.2017.01.050. PubMed PMID: 28162249.

52. Pasparakis C, Mager EM, Stieglitz JD, Benetti D, Grosell M. Effects of Deepwater Horizon crude oil exposure, temperature and developmental stage on oxygen consumption of embryonic and larval mahi-mahi (Coryphaena hippurus). Aquat Toxicol. 2016;181:113–23. Epub 2016/11/10. doi: 10.1016/j.aquatox.2016.10.022. PubMed PMID: 27829195.

53. Serrano XM, Miller MW, Hendee JC, Jensen BA, Gapayao JZ, Pasparakis C, et al. Effects of thermal stress and nitrate enrichment on the larval performance of two Caribbean reef corals. Coral Reefs. 2017;37(1):173–82. doi: 10.1007/s00338-017-1645-y.

54. Yashchenko V, Fossen EI, Kielland ON, Einum S. Negative relationships between population density and metabolic rates are not general. J Anim Ecol. 2016;85(4):1070–7. Epub 2016/03/13. doi: 10.1111/1365-2656.12515. PubMed PMID: 26970102.

55. Apprill A, McNally S, Parsons R, Weber L. Minor revision to V4 region SSU rRNA 806R gene primer greatly increases detection of SAR11 bacterioplankton. Aquatic Microbial Ecology. 2015;75(2):129–37. doi: 10.3354/ame01753.

56. Kuznetsova A, Brockhoff PB, Christensen RHB. lmerTest Package: Tests in Linear Mixed Effects Models. Journal of Statistical Software. 2017;82(13). doi: 10.18637/jss.v082.i13.

57. Ludecke MD. Package sjPlot: Data Visualization for Statistics in Social Science. 2022.

58. Palacio-Castro AM, Serrano XM. Data and code for “Sediment source and concentration influence larval performance of the threatened coral Orbicella faveolata” submitted to PlosOne (Version v1) [Data set]. Zenodo. 2023. doi: 10.5281/zenodo.8022069.

59. Bolyen E, Rideout JR, Dillon MR, Bokulich NA, Abnet CC, Al-Ghalith GA, et al. Reproducible, interactive, scalable and extensible microbiome data science using QIIME 2. Nat Biotechnol. 2019;37(8):852–7. Epub 2019/07/26. doi: 10.1038/s41587-019-0209-9. PubMed PMID: 31341288; PubMed Central PMCID: PMCPMC7015180.

60. Martin M. Cutadapt removes adapter sequences from high-throughput sequencing reads. EMBnet journal. 2011;17(1):10–2.

61. Callahan BJ, McMurdie PJ, Rosen MJ, Han AW, Johnson AJ, Holmes SP. DADA2: High-resolution sample inference from Illumina amplicon data. Nat Methods. 2016;13(7):581–3. Epub 2016/05/24. doi: 10.1038/nmeth.3869. PubMed PMID: 27214047; PubMed Central PMCID: PMCPMC4927377.

62. Bokulich NA, Kaehler BD, Rideout JR, Dillon M, Bolyen E, Knight R, et al. Optimizing taxonomic classification of marker-gene amplicon sequences with QIIME 2’s q2-feature-classifier plugin. Microbiome. 2018;6(1):90. Epub 2018/05/19. doi: 10.1186/s40168-018-0470-z. PubMed PMID: 29773078; PubMed Central PMCID: PMCPMC5956843 The authors declare that they have no competing interests. PUBLISHER’S NOTE: Springer Nature remains neutral with regard to jurisdictional claims in published maps and institutional affiliations.

63. Lin H, Peddada SD. Analysis of compositions of microbiomes with bias correction. Nat Commun. 2020;11(1):3514. Epub 2020/07/16. doi: 10.1038/s41467-020-17041-7. PubMed PMID: 32665548; PubMed Central PMCID: PMCPMC7360769.

64. Hodgson G. Sediment and the settlement of larvae of the reef coral Pocillopora damicornis. Coral Reefs. 1990;9:41–3.

65. Krausfeldt LE, Lopez JV, Bilodeau CM, Won Lee H, Casali SL. Change and stasis of distinct sediment microbiomes across Port Everglades Inlet (PEI) and the adjacent coral reefs. PeerJ. 2023;11:e14288. Epub 2023/01/20. doi: 10.7717/peerj.14288. PubMed PMID: 36655050; PubMed Central PMCID: PMCPMC9841897.

66. Klaus JS, Janse I, Fouke BW. Coral black band disease microbial communities and genotypic variability of the dominant cyanobacteria (CD1C11). Bulletin of Marine Science. 2011;87(4):795.

67. Waite DW, Chuvochina M, Pelikan C, Parks DH, Yilmaz P, Wagner M, et al. Proposal to reclassify the proteobacterial classes Deltaproteobacteria and Oligoflexia, and the phylum Thermodesulfobacteria into four phyla reflecting major functional capabilities. Int J Syst Evol Microbiol. 2020;70(11):5972–6016. Epub 2020/11/06. doi: 10.1099/ijsem.0.004213. PubMed PMID: 33151140.

68. Nie S, Zhang Z, Mo S, Li J, He S, Kashif M, et al. Desulfobacterales stimulates nitrate reduction in the mangrove ecosystem of a subtropical gulf. Sci Total Environ. 2021;769:144562. Epub 2021/01/19. doi: 10.1016/j.scitotenv.2020.144562. PubMed PMID: 33460836.

69. Clausing RJ, Annunziata C, Baker G, Lee C, Bittick SJ, Fong P. Effects of sediment depth on algal turf height are mediated by interactions with fish herbivory on a fringing reef. Marine Ecology Progress Series. 2014;517:121–9. doi: 10.3354/meps11029.

70. Dobbelaere T, Muller EM, Gramer LJ, Holstein DM, Hanert E. Coupled Epidemio-Hydrodynamic Modeling to Understand the Spread of a Deadly Coral Disease in Florida. Frontiers in Marine Science. 2020;7. doi: 10.3389/fmars.2020.591881.

71. Precht WF, Gintert BE, Robbart ML, Fura R, van Woesik R. Unprecedented Disease-Related Coral Mortality in Southeastern Florida. Sci Rep. 2016;6:31374. Epub 2016/08/11. doi: 10.1038/srep31374. PubMed PMID: 27506875; PubMed Central PMCID: PMCPMC4979204.

72. O’Connell L, Gao S, McCorquodale D, Fleisher J, Lopez JV. Fine grained compositional analysis of Port Everglades Inlet microbiome using high throughput DNA sequencing. PeerJ. 2018;6:e4671. Epub 2018/05/16. doi: 10.7717/peerj.4671. PubMed PMID: 29761039; PubMed Central PMCID: PMCPMC5947159.

73. Abdel-Salam HA, Porter JW, Hatcher B, editors. Physiological effects of sediment rejection on photosynthesis and respiration in three Caribbean reef corals. Proc 6th int coral Reef Symp; 1988.

74. Okubo N, Yamamoto HH, Nakaya F, Okaji K. Oxygen consumption of a single embryo/planula in the reef-building coral Acropora intermedia. Marine Ecology Progress Series. 2008;366:305–9. doi: 10.3354/meps07562.

75. Brainard R, Birkeland C, Eakin CM, McElhany P, Miller MW, Patterson M, et al. Status review report of 82 candidate coral species petitioned under the US Endangered Species Act: Citeseer; 2011.

76. Hughes TP, Tanner JE. Recruitment failure, life histories, and long-term decline of Caribbean corals. Ecology. 2000;81(8):2250–63.

77. Protection AE. Technical Guidance: Environmental Impact Assessment of Marine Dredging Proposals. Environmental Protection Authority, Perth, Western Australia. 2016.

78. Szmant AM. Reproductive ecology of Caribbean reef corals. Coral reefs. 1986;5:43–53.

79. Szmant AM. Sexual reproduction by the Caribbean reef corals Montastrea annularis and M. cavernosa. Mar Ecol Prog Ser. 1991;7(4):13–25.

