## Supplementary figures and images for "Sediment source and dose influence the larval performance of the threatened coral *Orbicella faveolata*"

### S1 Figure

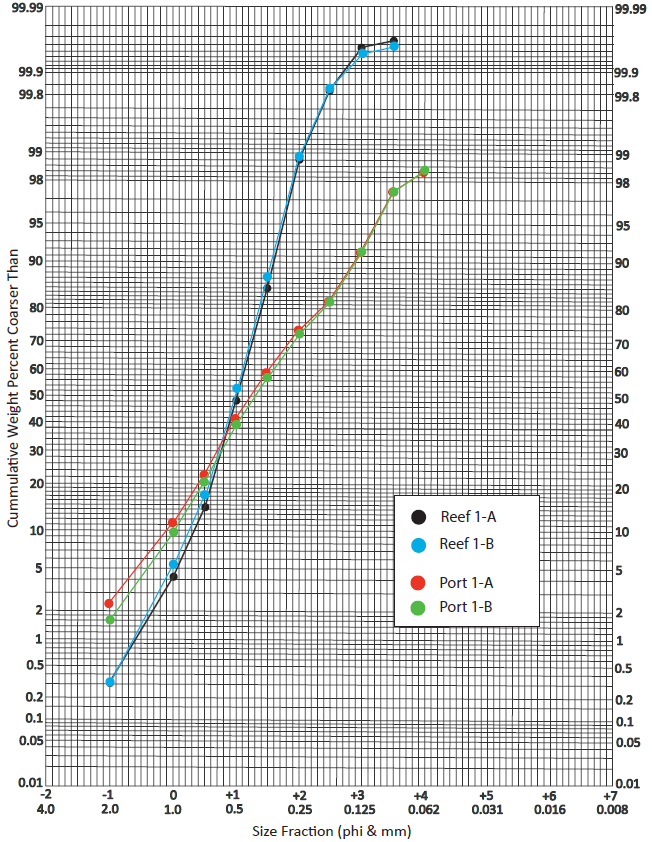

### S2 Figure

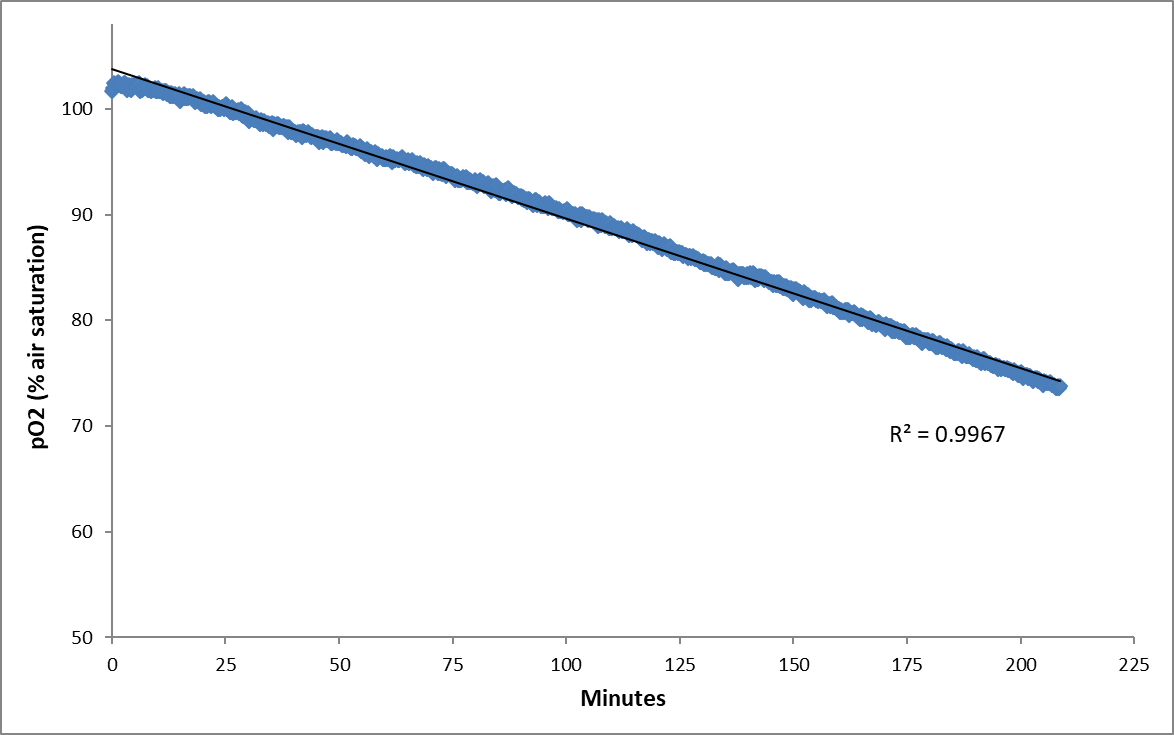

### S3 Figure

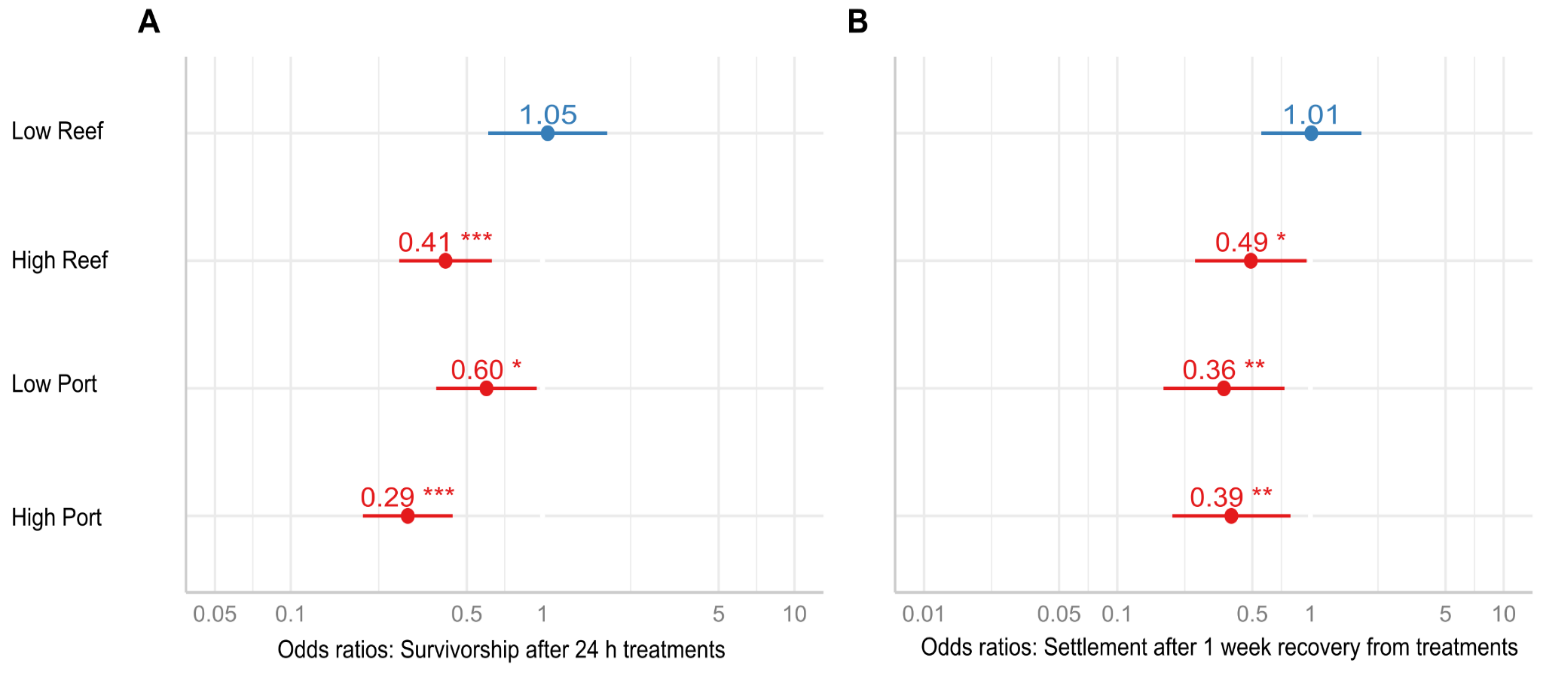

### S4 Figure

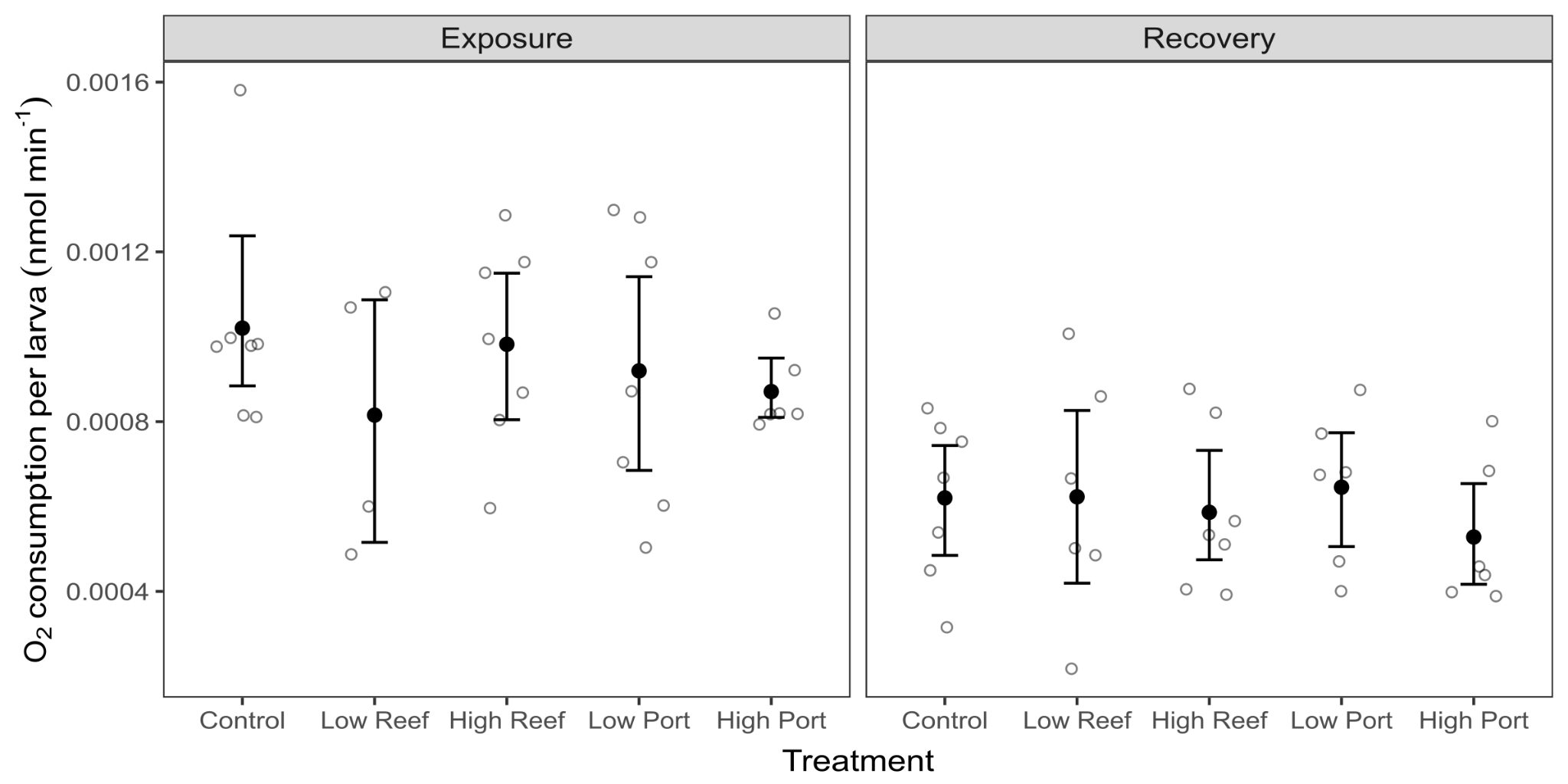
