## Supplementary material for "Sediment source and dose influence the larval performance of the threatened coral *Orbicella faveolata*": S1 Table

| **Sample** | **Port 1-A** | **Port 1-B** | **Reef 1-A** | **Reef 1-B** |
| --- | --- | --- | --- | --- |
| **Graphic Mean** | 1.37 phi  (0.39 mm) | 1.38 phi  (0.38 mm) | 1.0 phi  (0.50 mm) | 0.96 phi  (0.52 mm) |
| **Graphic Standard Deviation**  **(68% of distribution)** | 1.18 phi | 1.13 phi | 0.45 phi | 0.49 phi |
| **Inclusive Graphic**  **Standard Deviation (sorting)** | 1.14 phi  (poorly sorted) | 1.11 phi  (poorly sorted) | 0.49 phi  (well sorted) | 0.50 phi  (well sorted) |
| **Percent mud (<63 μm)** | 1.2 | 1.3 | 0.0 | 0.0 |
| **Percent sand (63 μm – 2 mm)** | 97.4 | 98.1 | 99.7 | 99.7 |
| **Percent gravel**  **(> 2 mm)** | 2.3 | 1.6 | 0.3 | 0.3 |
