## Supplementary material for "Sediment source and dose influence the larval performance of the threatened coral *Orbicella faveolata*": S2 Table

| MODEL | **DF** | **Deviance** | **Residual DF** | **Residual Deviance** |
| --- | --- | --- | --- | --- |
| **Survivorship model (24 h):** chi-square = 1.83x10-12 | | | | |
| NULL | NA | NA | 46 | 241.35 |
| Treatment | 4 | 60.96 | 42 | 180.40 |
| **Settlement model (1 week):** chi-square = 0.0021 | | | | |
| NULL | NA | NA | 43 | 54.53 |
| Treatment | 4 | 16.74 | 39 | 37.79 |
