## Supplementary material for "Sediment source and dose influence the larval performance of the threatened coral *Orbicella faveolata*": S3 Table

|  | A) **Larval survival** | | | | B) **Larval settlement** | | | |
| --- | --- | --- | --- | --- | --- | --- | --- | --- |
|  | **Coefficient estimate** | **Std. Error** | **z value** | **Pr(>\|z\|)** | **Coefficient estimate** | **Std. Error** | **z value** | **Pr(>\|z\|)** |
| **Intercept** | 2.7183 | 0.26015 | 10.449 | < 2e-16 *** | -1.2961 | 0.20956 | -6.185 | 6.21e-10 *** |
| **Low Reef** | 0.0466 | 0.27783 | 0.168 | 0.8667 | 0.0109 | 0.30499 | 0.036 | 0.97137 |
| **High Reef** | -0.8878 | 0.21637 | -4.103 | 4.08e-05 *** | -1.0311 | 0.30499 | -2.802 | 0.00507 ** |
| **Low Port** | -0.5134 | 0.23426 | -2.191 | 0.0284 * | -0.7104 | 0.33885 | -2.096 | 0.03604 * |
| **High Port** | -1.2327 | 0.2095 | -5.884 | 4.01e-09 *** | -0.9429 | 0.35922 | -2.625 | 0.00867 ** |
