## Supplementary material for "Sediment source and dose influence the larval performance of the threatened coral *Orbicella faveolata*": S4 Table

|  | **Sum Sq** | **Mean Sq** | **DF** | **DenDF** | **F value** | **Pr(>F)** |
| --- | --- | --- | --- | --- | --- | --- |
| Treatment | 7.57e-08 | 1.89e-08 | 4 | 41.18 | 0.63 | 0.64 |
| Timepoint | 4.74e-07 | 4.74e-07 | 1 | 1.94 | 15.83 | 0.06 |
| Treatment:Timepoint | 6.81e-08 | 1.70e-08 | 4 | 33.56 | 0.57 | 0.69 |
